# CSDE1 regulates miR-20a-5p/ TMBIM6 axis in melanoma

**DOI:** 10.1101/2025.01.07.631786

**Authors:** Sushmitha Ramakrishna, Tanit Guitart, Fatima Gebauer, Pavan Kumar Kakumani

**Affiliations:** Department of Biochemistry, Memorial University of Newfoundland, 45 Arctic Ave., St. John’s, NL Canada A1C 5S7; Gene Regulation, Stem Cells and Cancer Programme, Centre for Genomic Regulation (CRG), The Barcelona Institute of Science and Technology, 08003 Barcelona, Spain; Universitat Pompeu Fabra (UPF), 08003 Barcelona, Spain

**Keywords:** CSDE1, AGO2, miR-20a-5p, TMBIM6, gene silencing, melanoma

## Abstract

RNA-binding proteins (RBPs) and microRNAs (miRNAs) play crucial roles in regulating gene expression at the post-transcriptional level in tumorigenesis. They primarily target the 3′UTRs of mRNAs to control their translation and stability. However, their co-regulatory effects on specific mRNAs in the pathogenesis of particular cancers are yet to be fully explored. CSDE1 is an RBP that promotes melanoma metastasis, and the mechanisms underlying its function in melanoma development are yet to be fully understood. In the current study, we report that CSDE1 enhances TMBIM6 protein expression without altering its mRNA levels in melanoma cells, indicating post-transcriptional regulation. CSDE1 and AGO2 competitively bind to TMBIM6 mRNA, and we identify miR-20-5p, which represses TMBIM6 expression, regulates the binding of CSDE1 to TMBIM6 mRNA. Further, the RNA-binding mutant of CSDE1 foregoes its competitive binding to the mRNA, allowing AGO2-mediated silencing of TMBIM6 expression. Our study highlights the pivotal role of CSDE1 in regulating miR-20a-5p function and the expression of TMBIM6 in melanoma cells, thus unveiling the potential of therapeutic strategies targeting this regulatory pathway in treating malignant skin cancers.

## INTRODUCTION

Melanoma is the most aggressive form of skin cancer, and its incidence, as well as associated mortality, continues to increase worldwide (Schadendorf et al. 2015). According to the 2020 statistics published by Globocon, an estimated 57,000 individuals worldwide lost their lives to melanoma(Saginala et al. 2021). Understanding the mechanisms of genes responsible for melanoma development can help better treat this malignant form of skin cancer. MicroRNAs (miRNAs) are a special class of small non-coding RNAs that function as either oncogenes or tumor suppressors, and their altered expression impacts well-known oncogenic pathways such as the PI3K/AKT and RAS/MAPK cascade in melanoma tumorigenesis (Xu et al. 2012; Aksenenko et al. 2019). miRNAs are 21-23 nucleotides (nt) long and regulate gene expression at the post-transcriptional level, by binding to complementary sites in the 3’ untranslated regions (UTR) of messenger RNAs (mRNAs) (Kakumani et al. 2021, 2020; O’Brien et al. 2018). miRNAs are transcribed by RNA polymerase II in the nucleus, where these transcripts, referred to, as primary miRNAs (pri-miRNAs) are processed by the microprocessor complex, consisting of Drosha-DGCR8, to generate precursor miRNAs (pre-miRNAs), which are transported to the cytoplasm for subsequent maturation by Dicer complex into miRNAs (Hynes and Kakumani 2024). The mature miRNAs form miRNA-induced silencing complex (miRISC), of which AGO proteins are the essential components (Gebert and MacRae 2019). Initially, the miRNA duplex is loaded onto AGO, and the passenger strand is released, guiding the miRNA-AGO complex to complementary sites in mRNA 3’UTRs. Once the miRNA-mRNA interaction is initiated, AGO recruits GW182 (known as TNRC6 in humans) and CCR4-NOT complexes to facilitate translational repression and/or deadenylation, followed by the decay of targeted mRNAs (Jonas and Izaurralde 2015). Here, the target abundance, the number and accessibility of miRNA-binding sites, or the binding of specific RNA-binding proteins (RBPs) to target 3′UTRs play critical roles in miRNA-AGO binding to the targets and, thereby, miRNA silencing efficiency (Kakumani et al. 2021).

RBPs are essential players in mRNA metabolism, which regulate splicing, transport, translation, and degradation (Hentze et al. 2018). Interestingly, about half of the RBPs identified thus far bind mRNAs and manifest their function by regulating the fate of target mRNAs. Altered expression, localization, and post-translational modification of RBPs contribute to tumorigenesis by increasing the expression of oncogenes or decreasing the expression of tumor suppressor genes (Pereira et al. 2017; Neelamraju et al. 2018). RBPs recognize common mRNA features such as the 3′ poly(A) tail and the sequence motifs or secondary structures present in mRNA 3′UTRs (Hentze et al. 2018). The 3′UTRs of mammalian mRNAs can be as long as 10 kilobases and are bound by different miRNAs and RBPs (Jiang and Coller 2012). While miRNAs can only bind their target mRNAs in association with AGOs, RBPs directly bind their targets either as single entities or in complex with other RBPs to control mRNA metabolism. miRNA binding in 3′UTRs of mRNAs facilitates an intricate network of interactions between miRNA-AGO and RBPs, thus determining the fate of overlapping targets (Kakumani 2022). Examples involving RBPs and miRNAs postulate several cooperative and antagonistic interaction modes. For instance, HuR antagonizes miR-122-mediated repression of CAT-1 mRNA but promotes miRISC recruitment to let-7 miRNA site on c-Myc 3′UTR (Ahuja et al. 2016; Kim et al. 2009; Bhattacharyya et al. 2006). Besides, transcriptome-wide studies provided evidence for antagonistic and cooperative AGO2-RBP interactions in the case of HuR, Pumilio, and FAM120A toward mRNA regulation (Kedde et al. 2010; Sternburg et al. 2018; Kelly et al. 2019; Li et al. 2018; Kundu et al. 2012). However, the effects of RBPs on the functioning of miRNA-AGO complex relevant to a specific biological or pathological context are yet to be fully documented, especially in the case of melanoma.

The RBP, Cold shock domain (CSD) containing protein E1 (CSDE1), first described as Unr (upstream of N-ras), plays a vital role in translational reprogramming, mainly expressed in the cell cytoplasm (Moore et al. 2018; Guo et al. 2020; Kim et al. 2022). CSDE1 contains at least five canonical CSDs (Hollmann et al. 2020). The N-terminal CSDs, namely CSD1 and CSD2, are crucial for their RNA-binding ability, whereas the C-terminal CSDs are involved in protein-protein interactions (Ciocia et al. 2024). In *Drosophila melanogaster*, CSDE1 is part of a translational repressor complex assembled at the 3′UTR of male-specific-lethal-2 mRNA and is critical for proper regulation of X-chromosome dosage compensation (Abaza et al. 2006; Duncan et al. 2006). CSDE1 was shown to interact with PABP to control the stability of c-fos mRNA and with AUF1 to regulate PTH mRNA (Dinur et al. 2006). Further, CSDE1 regulates the stability and translation of FABP7 and Vimentin (VIM) mRNAs in human embryonic stem cells (ESCs), while it controls GATA6 post-transcriptionally in mouse ESCs (Ju Lee et al. 2017; Elatmani et al. 2011). In melanoma, CSDE1 acts as an oncogene and promotes the translation elongation of critical epithelial-to-mesenchymal transition (EMT) markers, namely VIM and RAC1, to enhance invasion and metastasis (Wurth et al. 2016). In our previous study, we showed that CSDE1 interacts with numerous miRISCs in *Drosophila* embryo extracts, mouse, and human cell lines and further associates with DCP1-DCP2 decapping complex to render target mRNAs susceptible to miRNA-guided gene silencing (Kakumani et al. 2020). Subsequently, we discovered that CSDE1 and AGO2 have overlapping binding sites on the 3’UTR of numerous target mRNAs, especially prostate transmembrane protein, androgen-induced 1 (PMEPA1) mRNA, and CSDE1 counteracts AGO2/miRISC-guided translational repression of PMEPA1 to promote melanoma tumorigenesis (Kakumani et al. 2021). However, as CSDE1 binds a large target mRNA repertoire in melanoma (Wurth et al. 2016), the molecular details on its contributions to specific target mRNA regulation are still incomplete.

In the present study, we aim to expand the current understanding of the regulatory effects of CSDE1 in miRISC-mediated repression of target mRNAs, commonly bound by both CSDE1 and AGO2, in melanoma. In particular, we focused on the relationship between CSDE1 and the miR-20a-5p/AGO2 binding to, and the regulation of transmembrane BAX inhibitor motif containing 6 (TMBIM6) mRNA in melanoma cells. We demonstrated that the miRNA pathway, particularly miR-20a-5p suppresses TMBIM6 expression in melanoma cells. Whereas CSDE1 hinders AGO2 binding to TMBIM6 mRNA, thereby promoting TMBIM6 protein expression, with implications in melanoma tumorigenesis.

## RESULTS

### CSDE1 promotes the expression of TMBIM6 in melanoma

CSDE1 binds a total of 396 target mRNAs in melanoma cells that are linked to cancer development (Wurth et al. 2016). In our previous study, we analyzed the CSDE1 binding pattern on the target mRNAs and identified multiple candidates which are bound by both CSDE1 and AGO2 on the 3’UTRs, including TMBIM6 and CDK6 (Supplemental File 1) (Kakumani et al. 2021). To study the relation between CSDE1 and the identified targets, we measured the effects of CSDE1 depletion and overexpression on CDK6 and TMBIM6 levels in metastatic (SK-Mel-147) and non-metastatic (UACC-62) melanoma cells. We observed that the loss of CSDE1 resulted in the reduction of TMBIM6 protein levels in both the cell types, while it did not affect CDK6 protein expression (Fig. 1A, 1C (Top)). Next, we quantified the mRNA levels of both TMBIM6 and CDK6 under the same conditions and observed that there was no significant difference in their transcript levels (Fig. 1A, 1C (bottom)). In corollary, we measured TMBIM6 and CDK6 expression levels in the melanoma cells transiently expressing FLAG-tagged CSDE1 and observed that TMBIM6 protein was upregulated but noticed no significant change in the expression levels of CDK6 (Fig. 1B, 1D (top)). Additionally, we quantified the mRNA levels of both TMBIM6 and CDK6 in both SK-Mel-147 and UACC-62 cells under the same conditions and observed no altered expression in cells transient overexpressing CSDE1, compared to the vector control (Fig. 1B, 1D (bottom)). Following, we tested the possibility of CSDE1 influencing TMBIM6 protein stability. We treated the CSDE1 knockdown melanoma cells with MG132, a proteasome inhibitor, and measured TMBIM6 protein levels. The TMBIM6 expression was downregulated in CSDE1 knockdown conditions relative to the control. However, it was unaltered between the conditions, treated or not, with MG132, indicating the effects of CSDE1 on TMBIM6 are independent of the proteasomal degradation pathway (Fig. S1A). Further, we observed a correlation between TMBIM6 and CSDE1 protein expression in multiple melanoma cell lines and no expression of both the proteins in non-tumoral melanocytes (Fig. S1B). Overall, our results show that CSDE1 positively regulates TMBIM6 expression in melanoma.

**Figure 1:**
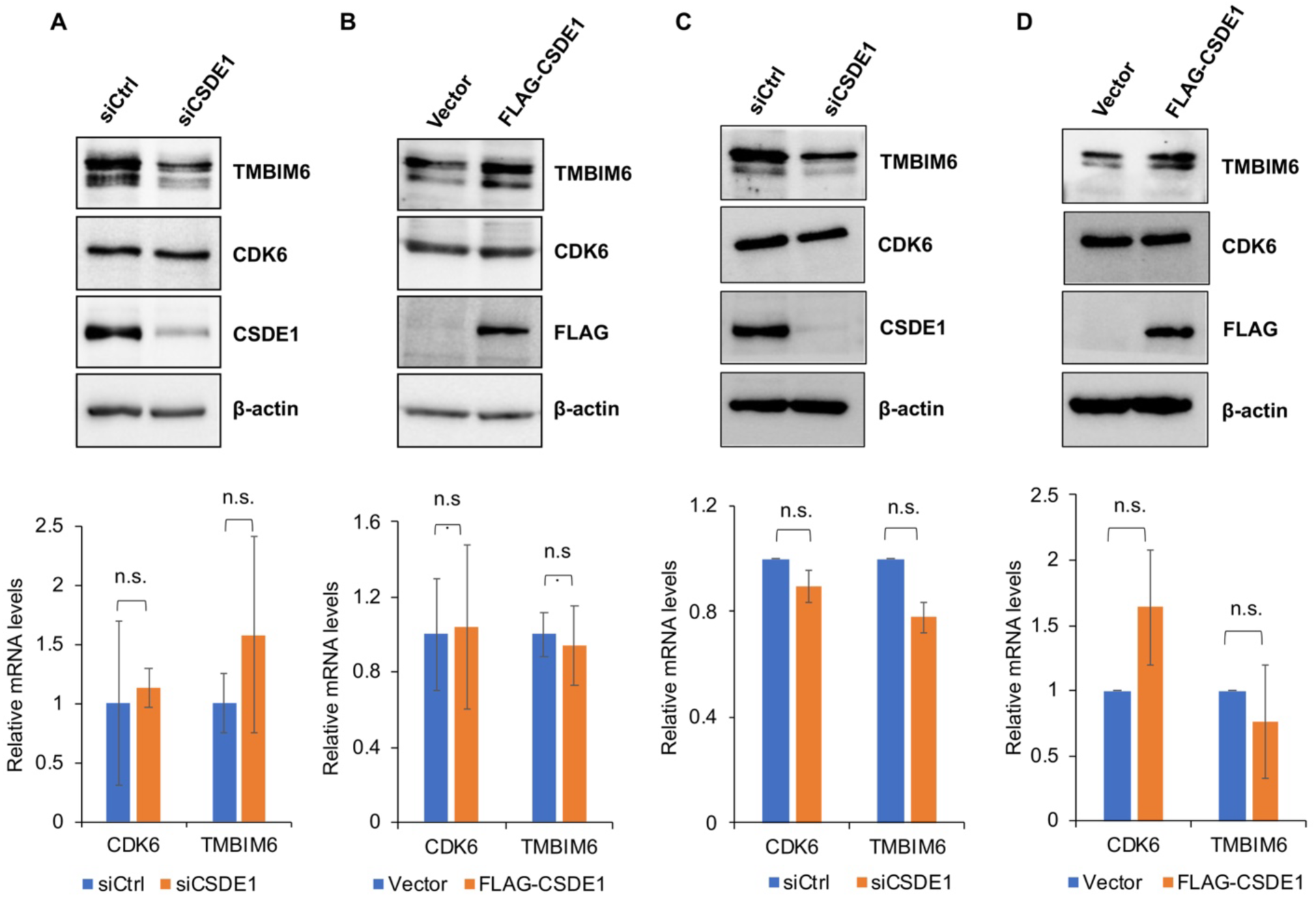
CSDE1 promotes the expression of TMBIM6 in melanoma cells. **(A)** (Top) Western blots showing the expression of protein targets under control and CSDE1 knockdown conditions in SK-Mel-147 cells. (Bottom) Relative CDK6 and TMBIM6 mRNA levels in total RNA from control and CSDE1 depleted SK-Mel-147 cells. RT-qPCR was performed to quantify the mRNA levels. **(B)** Western blot analysis of proteins indicated in SK-Mel-147 cells with transiently expressed FLAG-tagged CSDE1 or control vector. (Bottom) Relative CDK6 and TMBIM6 mRNA levels in total RNA from vector control and wild-type FLAG-CSDE1. **(C)** (Top) Western blots probed for target proteins under control and CSDE1 knockdown conditions in UACC-62 cells. (Bottom) Relative CDK6 and TMBIM6 mRNA levels in total RNA from control and CSDE1 depleted UACC-62 cells. RT-qPCR was performed to quantify the mRNA levels. **(D)** Western blot analysis of target protein expression in UACC-62 cells with transiently expressed FLG-tagged CSDE1 or the control vector. (Bottom) Relative CDK6 and TMBIM6 mRNA levels in total RNA from vector control and the wild-type FLAG-CSDE1. Data in (A-D, Bottom) are presented as mean ± SD. (n.s., non-significant, p>0.05; n=3; two tailed t-test). For both western blotting and RT-qPCR analysis, β-actin served as the loading control and the reference gene, respectively.

### miR-20a-5p negatively regulates TMBIM6 expression

TMBIM6 is overexpressed in numerous cancer types, including liver, prostate, lung, and breast cancers, and the overall survival of patients is dependent on TMBIM6 expression, especially in skin cutaneous melanoma (Kim et al. 2020). Since miRNAs regulate numerous hallmarks of melanoma (Gajos-Michniewicz and Czyz 2019; Poniewierska-Baran et al. 2022), we tested whether the miRNA pathway influences TMBIM6 expression in melanoma cells. We knocked down two essential miRNA pathway components, namely, Dicer and AGO2, in SK-Mel-147 cells and measured the expression levels of TMBIM6. We observed increased expression of TMBIM6 protein in both Dicer and AGO2 knockdown conditions relative to the control, indicating that the miRNA pathway negatively regulates TMBIM6 expression in metastatic melanoma cells, while it had no influence on CSDE1 levels (Fig. 2A-B). Next, to identify miRNAs that silence TMBIM6 expression, we initially identified putative miRNAs that target the 3’UTR of TMBIM6 using the online tool TargetScan (Agarwal et al. 2015). Based on the minimum free energy of binding between the miRNA seed region and its complementary site in the 3’UTR, we shortlisted the candidate miRNAs, miR-19a-3p, miR-20a-5p, miR-27a-3p, miR-125b-5p, miR-183-5p. Following, we transfected melanoma cells with complementary oligos (antagomiRs) against the candidate miRNAs and measured TMBIM6 expression. We observed an increased expression of TMBIM6 protein in the case of miR-19a-3p, miR-20a-5p and miR-27a-3p, relative to the control (Fig. 2C). However, in light of the upregulation of PMEPA1, another CSDE1 target (Kakumani et al. 2021), in the case of miR-19a-3p and miR-27a-3p, we chose to proceed with miR-20a-5p, to demonstrate the direct relation between miR-20a-5p and TMBIM6. Here, we transfected the melanoma cells with mimics of miR-20a-5p and noted a decrease in the expression of TMBIM6 protein relative to the control (Fig. 2D). Subsequently, to determine the translational repression of TMBIM6 is mediated by direct binding of miR-20a-5p to its 3’UTR, at first, we carried out luciferase reporter assays by using constructs with and without TMBIM6 3’UTR. As seen in Fig. 2E, the relative luciferase activity was reduced in the presence of TMBIM6 3’UTR, compared to the empty vector, suggesting that variations in TMBIM6 expression are related to its 3’UTR. Following, we performed the reporter assays using constructs with point mutations in the miR-20a-5p target sites in the TMBIM6 3’UTR (Fig. 2F, left). We observed enhanced luciferase activity in the case of the mutant, relative to the wild-type 3’UTR (Fig. 2F, right), revealing that miR-20a-5p binding is, in fact, necessary for translational repression of TMBIM6.

**Figure 2:**
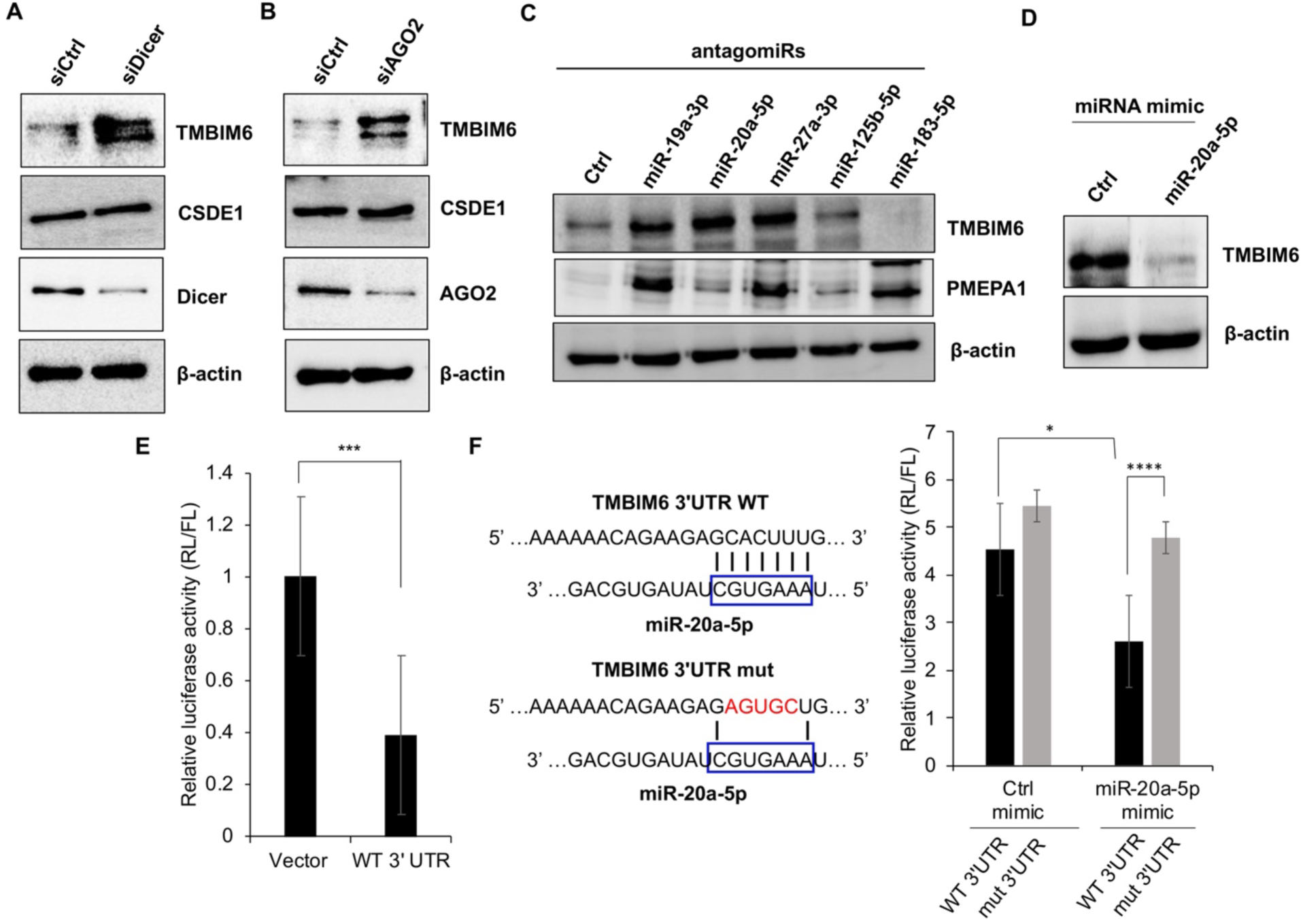
The miRNA pathway regulates TMBIM6 expression. **(A)** TMBIM6 expression analysis by western blotting using the lysate from SK-Mel-147 cells under control and Dicer knockdown conditions. **(B)** Detection of TMBIM6 by western blotting in lysates prepared from SK-Mel-147 cells under control and AGO2 knockdown conditions. **(C)** Western blots showing the expression of indicated proteins in lysates prepared from SK-Mel-147 cells transfected with antagomiRs of the control or the indicated miRNAs. **(D)** TMBIM6 expression analysis by western blotting from SK-Mel-147 cell lysate transfected with mimics of the control or miR-20a-5p. **(E)** Relative luciferase levels (renilla/firefly) from the lysate of HEK293T cells transfected with the vector control or the TMBIM6 3’UTR reporter construct. **(F)** Left: The representation of miR-20a-5p base pairing with either TMBIM6 wild-type (WT) or mutated (mut) 3’UTR sequence. The nucleotides in the blue box represent the seed region of miR-20a-5p. Right: Relative luciferase levels (renilla/firefly) from the lysate of HEK293T cells co-transfected with miRNA mimics and the TMBIM6 3’UTR reporter constructs. Data in (E), and (F) are presented as mean ± SD. (*, p<0.05; ***, p<0.01; ****, p<0.001; n=3; two tailed t-test). β-actin served as the loading control.

### CSDE1 counters miR-20a-5p-mediated repression of TMBIM6

In melanoma, CSDE1 interacts with select miRISCs, including that of miR-20a-5p (Kakumani et al. 2021). Since CSDE1’s association with AGO2, an essential component of miRISC, depends on target mRNAs (Kakumani et al. 2021), we sought to verify whether CSDE1 interaction with miR-20a-5p/RISC is RNA-dependent. The miRISC was purified from SK-Mel-147 cell extracts using modified complementary oligonucleotides specific to miR-20a-5p, following which, the samples were treated with RNases and measured for the abundance of CSDE1. We observed an almost complete loss of CSDE1 in the case of samples treated with RNases (Fig. 3A), indicating CSDE1’s association with miR-20a-5p/RISC is RNA-dependent. Next, we studied whether miR-20a-5p regulates the binding of CSDE1 to its target mRNAs, namely TMBIM6. We transfected melanoma cells with antagomiRs specific to miR-20a-5p and measured the levels of TMBIM6 mRNA bound to CSDE1. As shown in Fig. 3C, there was a significant increase in the levels of TMBIM6 bound to CSDE1 in the case of samples transfected with antagomiRs against miR-20a-5p, relative to the control, while the mRNA levels were unaltered in the total RNA (Fig. 3B). Additionally, we observed an increased expression of TMBIM6 protein levels in the case of miR-20a-5p antagomiRs (Fig. 3D), indicating miR-20a-5p restricts the binding between CSDE1 and its target TMBIM6 mRNA. Following, to confirm the inverse relation between miR-20a-5p and CSDE1 in regulating the expression of TMBIM6, we transfected the melanoma cells with mimics of miR-20a-5p or the control, followed by transient overexpression of FLAG-CSDE1, and examined the expression levels of TMBIM6. As shown in Fig. 3E, the protein levels of TMBIM6 were reduced in the case of miR-20a-5p mimic relative to the control, and however, the protein levels were restored upon the overexpression of CSDE1, indicating CSDE1 hinders miR-20a-5p function to suppress TMBIM6 expression in melanoma cells.

**Figure 3:**
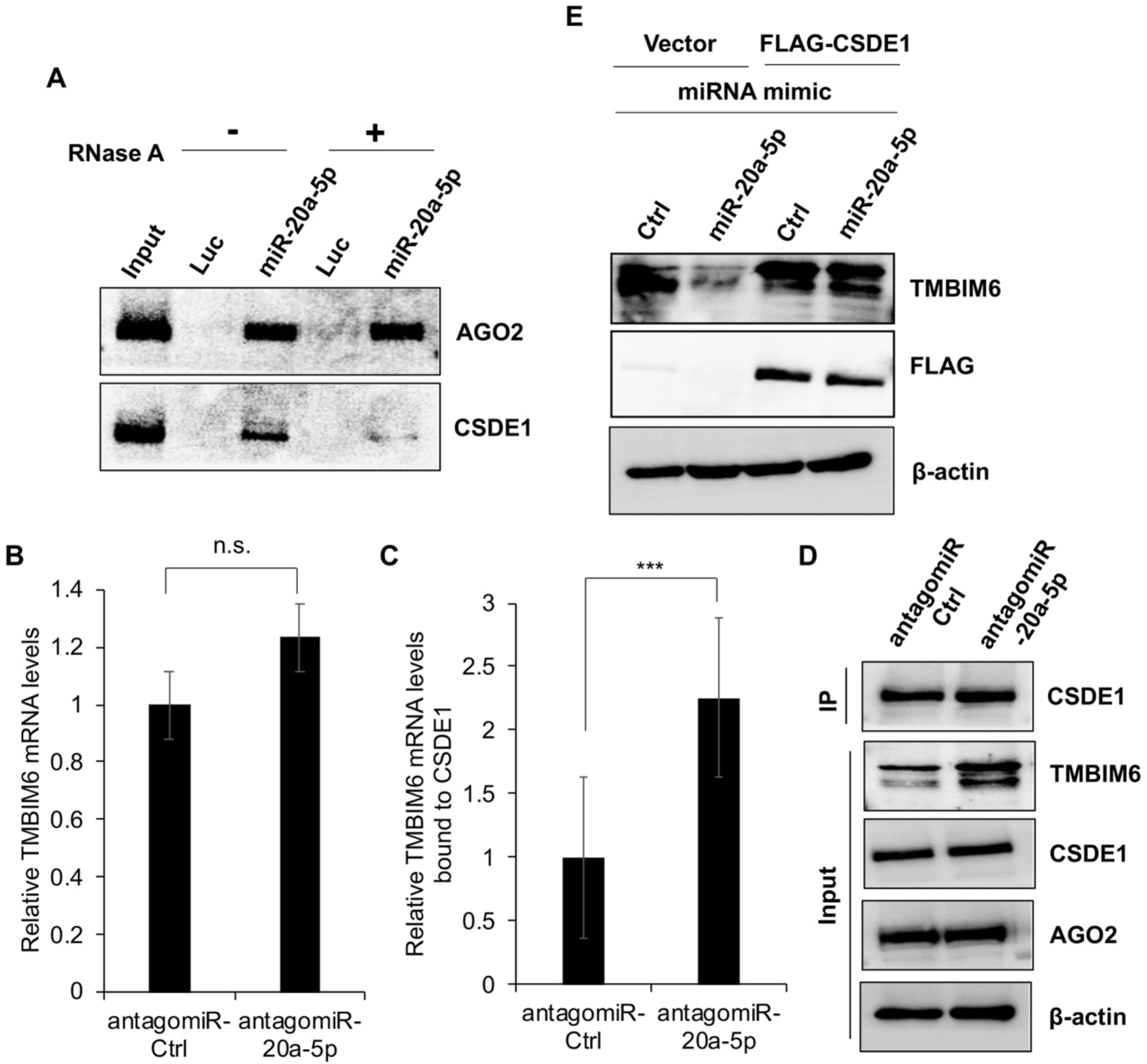
CSDE1 and miR-20a-5p oppose in regulating TMBIM6 expression. **(A)** Western blots confirming the expression of AGO2 and CSDE1 proteins in the samples from miR-20a-5p pull down using the lysate of SK-Mel-147 cells, before (-) and after (+) RNase treatment. **(B-C)** Relative TMBIM6 mRNA levels in total RNA **(B)** and that are bound by CSDE1 **(C)** in SK-Mel-147 cells transfected with the control or miR-20a-5p antagomiRs. **(D)** Western blot confirming the efficiency of CSDE1 immunoprecipitation from cells transfected with the control or miR-20a-5p antagomiRs. **(E)** Western blot analysis of TMBIM6 expression from SK-Mel-147 cell lysate transfected with miRNA mimics and co-transfected with FLAG-CSDE1. Data in (B-C) are presented as mean ± SD. (***, p<0.01; n.s., non-significant, p>0.05; n=3; two tailed t-test). The mRNA levels were quantified using dye-based RT-qPCR. β-actin served as a loading control and the reference gene for western blotting and dye-based RT-qPCR, respectively.

### CSDE1 and AGO2 compete to bind TMBIM6 mRNA

CSDE1 and AGO2 bind their target mRNAs on 3′UTRs to deliver downstream regulatory effects on gene expression (Wurth et al. 2016; Jonas and Izaurralde 2015; Gebert and MacRae 2019), and the target mRNA binding by AGO2 is critical in facilitating the interaction with CSDE1 in melanoma cells (Kakumani et al. 2021). Since both CSDE1 and AGO2 influence the expression of TMBIM6 (Fig. 1 and Fig. 2B), we hypothesized that they influence each other to fine-tune the expression of TIMBIM6. To test our hypothesis, we used CSDE1-depleted SK-Mel-147 (shCSDE1 SK-Mel-147) cells and performed AGO2 immunoprecipitation, followed by TMBIM6 mRNA and miR-20a-5p quantification. We observed an increase in the levels of TMBIM6 mRNA and miR-20a-5p bound to AGO2 under CSDE1 depleted conditions, compared to the control (Fig. 4B-C), while there was no significant difference in the mRNA levels in total RNA under the same conditions (Fig. 4A). However, the protein levels of TMBIM6 were downregulated (Fig. 4D), indicating that the enhanced binding of miR-20a-5p to AGO2 and the repression of TMBIM6 under CSDE1 depleted conditions is related to the binding of AGO2 to the target mRNA. Next, we examined whether AGO2 exerts similar effects on CSDE1 binding to the target mRNA. Here, we knocked down AGO2 in the melanoma cells and measured the amounts of TMBIM6 mRNA bound to CSDE1. We found that TMBIM6 mRNA levels bound by CSDE1 were higher in the case of AGO2 knockdown, compared to the control (Fig. 4F), while the mRNA and the miR-20a-5p levels in the total RNA were unaltered (Fig. 4E, 4G). Furthermore, we observed an increase in TMBIM6 protein levels under AGO2 knockdown conditions, an indication of the consequence of higher CSDE1 association with the TMBIM6 mRNA. Together, our findings demonstrate that CSDE1 and AGO2 compete to bind TMBIM6 mRNA to regulate its protein expression levels in melanoma.

**Figure 4:**
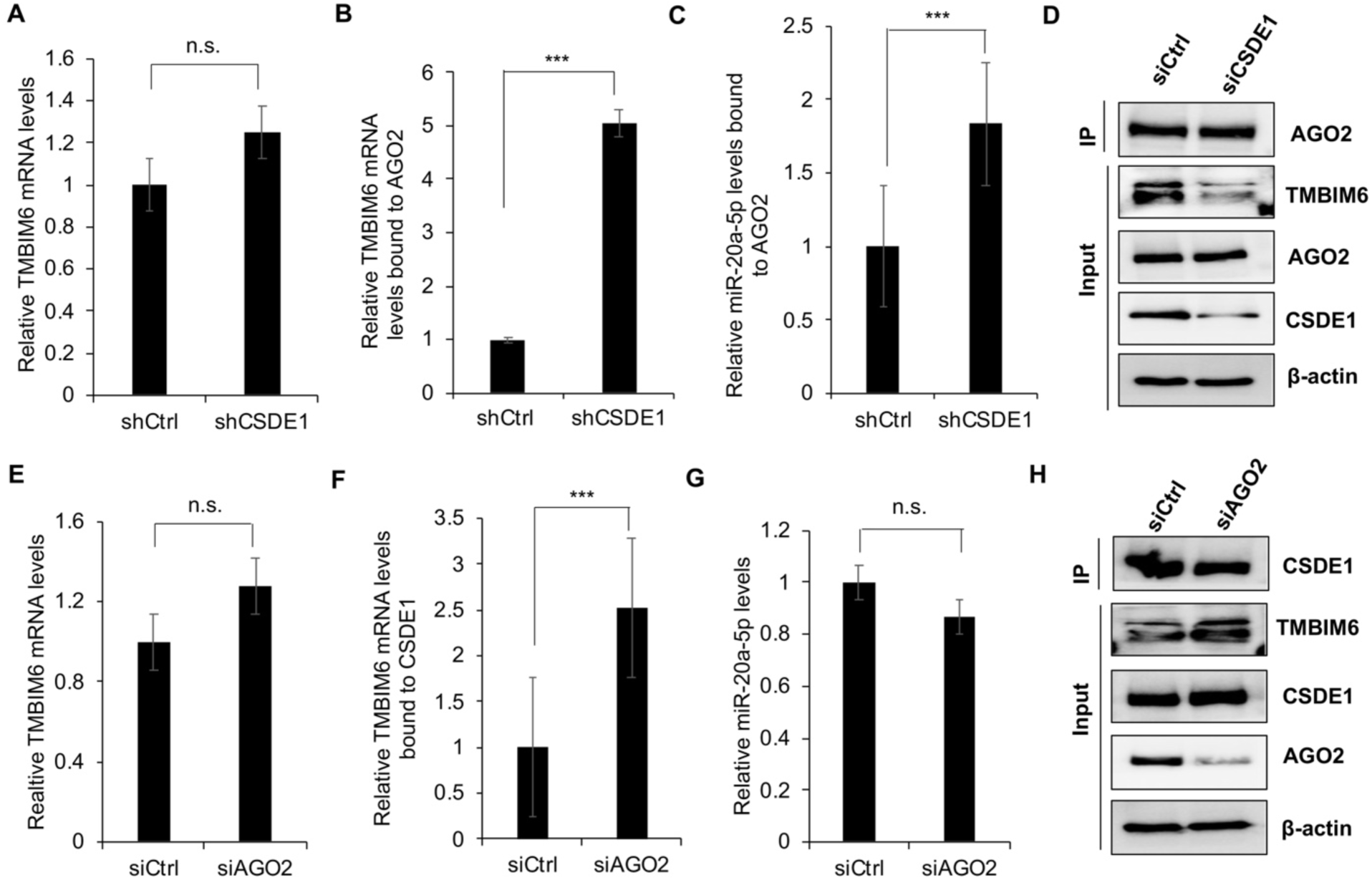
CSDE1 and AGO2 compete to bind TMBIM6 mRNA. (A-B) Relative mRNA levels of TMBIM6 in the total RNA **(A)** and that are bound to AGO2 **(B)** in shCtrl and shCSDE1 SK-Mel-147 cells. **(C)** Relative miR-20a-5p levels bound to AGO2 in shCtrl and shCSDE1 SK-Mel-147 cells. **(D)** Western blot confirming the depletion of CSDE1 in shCSDE1 cells relative to the shCtrl, and the efficiency of AGO2 immunoprecipitation between shCtrl and the shCSDE1 cell samples. **(E-F)** Relative mRNA levels of TMBIM6 in the total RNA **(E)** and that are bound by CSDE1 **(F)** in control and AGO2 knockdown SK-Mel-147 cells. **(G-H)** Relative miR-20a-5p levels in total RNA **(G)** and the western blots confirm the depletion of AGO2 and the efficiency of CSDE1 immunoprecipitation **(H)** in control and AGO2 knockdown SK-Mel-147 cells. Data in (A-C) and (E-G) are presented as mean ± SD. (***, p<0.01; n.s., non-significant, p>0.05; n=3; two tailed t-test). The mRNA and miRNA levels were quantified using dye-based RT-qPCR and TaqMan miRNA assays, respectively. β-actin served as a loading control and the reference gene for western blotting and dye-based RT-qPCR, respectively. For TaqMan miRNA assays, U6snRNA was used as a reference gene.

### The N-terminal CSD of CSDE1 controls AGO2 interaction with TMBIM6 mRNA

CSDE1 contains five canonical CSDs that serve to bind single-stranded nucleic acids and can function as protein-protein interaction modules (Mihailovich et al. 2010; Hollmann et al. 2020; Abaza and Gebauer 2008; Kamenska et al. 2016; Cornelis 2005). CSDE1 interacts with AGO2 via CSD1 (Kakumani et al. 2020, 2021) and interestingly, CSD1 is also responsible for the RNA-binding ability of CSDE1 (Mihailovich et al. 2010). To assess the contribution of CSDE1 binding to AGO2 and/or the RNA-binding ability, in the repression of TMBIM6, we generated a point mutant, Y37A, located in CSD1, which abrogates CSDE1’s capacity to bind RNAs. The schematics of the deletion and the point mutant of CSDE1 are illustrated in Fig. 5A. The recombinant FLAG-tagged clones were transiently expressed in CSDE1-depleted melanoma cells, and immunoprecipitation was performed using a FLAG antibody. As shown in Fig. 5B, the N-terminal deletion mutant ΔCSD1 and the point mutant Y37A showed reduced interaction with AGO2, compared to the wild-type CSDE1, indicating that CSD1, in particular, the RNA-binding capacity of CSDE1 is required for CSDE1 association with AGO2 in melanoma cells. Next, we examined the influence of the CSDE1 mutants on TMBIM6 expression under the same settings. We observed that the expression of wild-type CSDE1 restored the levels of TMBIM6 protein, while the deletion and the point mutants were unable to do so, compared to the vector control (Fig. 5C). Additionally, there was a reduction in the levels of TMBIM6 mRNA bound to the mutants ΔCSD1 and Y37A compared to the FLAG-tagged wild-type CSDE1 (Fig. 5E), which is consistent with the downregulation of TMBIM6 protein levels in the case of CSDE1 mutants relative to the wild-type, with no significant difference in TMBIM6 mRNA levels in the total RNA, across the samples (Fig. 5C-D). Further, we quantified TMBIM6 mRNA bound to AGO2 and observed that the target mRNA binding to AGO2 was higher in the case of both the mutants, compared to the samples expressing the wild-type CSDE1 (Fig. 5F-G), suggesting that increased TMBIM6 protein levels are associated with lower binding of AGO2 to TMBIM6 mRNA. Collectively, these results indicate that the interaction between CSDE1 and AGO2, facilitated by the RNA binding ability of CSDE1, via CSD1, is crucial for maintaining the expression of TMBIM6 in melanoma cells.

**Figure 5:**
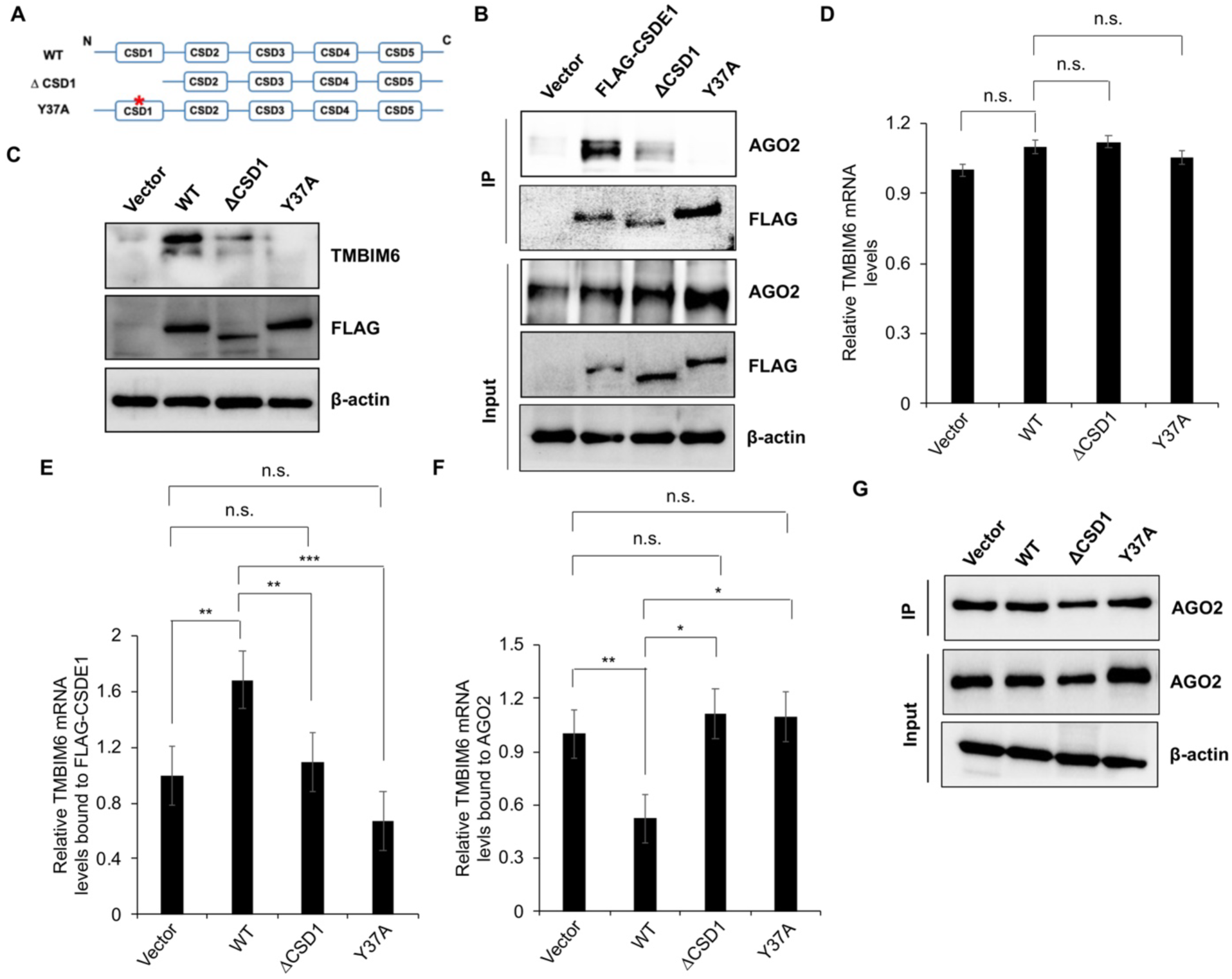
CSDE1 interaction with AGO2 regulates TMBIM6 expression. **(A)** Schematic representation of the wild-type CSDE1 and its deletion and point mutants used in the study. Only canonical Cold shock domains (CSDs) are illustrated. The non-canonical domains are not depicted. **(B)** Immunoprecipitation of FLAG-tagged CSDE1 and its deletion (ΔCSD1) and point (Y37A) mutants, using anti-FLAG antibody from the lysate of shCSDE1 SK-Mel-147 cells transiently expressing FLAG-CSDE1 and the mutants. AGO2 found in the immunoprecipitates was resolved by SDS-PAGE and detected by western blotting. **(C)** Western blots showing the expression of proteins indicated in the cell lysate of shCSDE1 SK-Mel-147 cells transiently expressing FLAG-CSDE1 and its mutants. **(D-F)** Relative TMBIM6 mRNA levels in the total RNA **(D)**, and that are bound to FLAG-CSDE1 **(E)** and AGO2 **(F),** in samples of shCSDE1 SK-Mel-147 cells transiently expressing the FLAG-CSDE1 and its mutants. Data in (D-F) are presented as mean ± SD. (*, p<0.05; **, p<0.03, ***, p<0.01; n.s., non-significant, p>0.05; n=3; two tailed t-test). **(G)** Western blot confirming the efficiency of AGO2 immunoprecipitation across the samples expressing FLAG-CSDE1 or its mutants. β-actin served as a loading control and the reference gene for western blotting and dye-based RT-qPCR, respectively.

## DISCUSSION

miRNAs primarily bind to 3’UTR of mRNA (Lai 2002) which is a hotspot for RBPs, in order to regulate the stability and translation of the target transcripts (Connerty et al. 2015). Our study delves into the regulatory dynamics of CSDE1 and AGO2 on TMBIM6 expression in melanoma cells, expanding our present understanding of the interplay between CSDE1 and the miRNA pathway.

In melanoma, CSDE1 interacts with specific miRNAs, and its association with AGO2 is RNA-dependent (Kakumani et al. 2021). CSDE1 and AGO2 share overlapping binding sites in the 3’UTR of target mRNAs, including PMEPA1 and TMBIM6 (Supplemental File 1) (Kakumani et al. 2021). In the current study, we demonstrate that CSDE1 promotes the expression of TMBIM6 at protein levels and their expression is concomitant across different melanoma cell types. Interestingly, the TMBIM6 mRNA levels were unaltered either in the case of knockdown or transient overexpression of CSDE1 (Fig. 1). Additionally, MG132 treatment of CSDE1 depleted cells showed no change in TMBIM6 protein expression (Fig. S1), suggesting an interplay between CSDE1 and AGO2 at the translation level to regulate TMBIM6 expression. These findings are consistent with the role of RBPs in modulating mRNA translation and stability, especially of CSDE1, which can alter the translation of target mRNAs without affecting their transcript levels (Kakumani 2022; Kakumani et al. 2021; Wurth et al. 2016). Although we observed the interaction between CSDE1 and the DCP1-DCP2 complex involved in mRNA decapping and decay (Kakumani et al. 2020), it is not uncommon for this RBP to behave differently depending on the biological and/or pathological context and the target mRNA (Wurth et al. 2016; Chang et al. 2004; Ciocia et al. 2024; Kakumani et al. 2020; Abaza et al. 2006). Similar observations have been made for other RBPs, such as Pumilio, which interferes with translation elongation, competes with eIF4E and even destabilizes target mRNAs depending on cellular background (Kaye et al. 2009; Friend et al. 2012; Bohn et al. 2018).

We observed that the core catalytic proteins of the miRNA pathway, namely Dicer and AGO2, negatively regulate TMBIM6 expression in melanoma cells (Fig. 2). Also, we identified miR-20a-5p as the negative regulator of TMBIM6 expression in melanoma cells. The miRNA, hsa-miR-20a-5p, is well conserved across animal species, and it belongs to the miR-17/92 cluster, with crucial roles in tumorigenesis, especially during invasion, metastasis, and inhibition of cell senescence. miR-20a-5p acts as an oncogene in breast, ovary, and hepatic carcinomas, and as a tumor suppressor in cervical and endometrial cancers (Huang et al. 2022). TMBIM6, as an anti-apoptotic protein, has a role in the development of cancers and was reported to be upregulated in breast, cervical, and prostate cancers (Lee et al. 2010). Since the overexpression of miR-20a-5p has been linked to melanoma, via the regulation of the TGF-β signalling pathway (Aksenenko et al. 2019), we believe that future studies investigating the downstream effects of CSDE1 on miR-20a-5p, and the negative regulation of TMBIM6 by miR-20a-5p, in manifesting tumorigenic capacities of melanoma cells will aid in an in-depth understanding of signalling cascades promoting melanoma development.

CSDE1 competes with AGO2 to bind the target mRNA to control TMBIM6 protein expression. The AGO2 knockdown and the presence of antagomiRs against miR-20a-5p enhanced the binding of TMBIM6 mRNA to CSDE1 (Fig. 3-4). Additionally, the loss of CSDE1 N-terminal CSD1 and its point mutant Y37A, which abrogates CSDE1’s ability to bind target RNAs and AGO2, enhances the binding of AGO2 to TMBIM6, leading to its repression (Fig. 5). Importantly, the CSD1 domain is required for CSDE1 RNA-binding as well as it is for AGO2 association, suggesting that target recognition and occupancy by CSDE1 determines the degree of recruitment of miRISC complex to specific mRNAs and their subsequent repression. These findings are consistent with our previous data from melanoma cells, in which CSDE1 promotes PMEPA1 expression by antagonizing AGO2/miRISC binding to the target mRNA. Similar mechanisms were also reported for other RBPs, including HuR and PUF (Friend et al. 2012; Kim et al. 2009; Kakumani 2022; Kakumani et al. 2021). Although our results expand on the number of studies demonstrating the miRNA machinery interaction with RBPs on specific mRNAs (Kedde et al. 2007; Young et al. 2012; Bottini et al. 2017; Xue et al. 2013; Kakumani 2022), in the future, we anticipate that transcriptome-wide analysis of CSDE1 and AGO2 binding patterns along melanoma progression would likely provide a comprehensive outlook on cooperative and antagonistic relationships between these two proteins, underlying the progression and severity of melanoma.

In summary, our study presents a mechanistic understanding of CSDE1 regulation of AGO2/miRISC recruitment to TMBIM6 mRNA (Fig. 6). We believe that future research should explore the broader implications of CSDE1 and AGO2 interactions with other common target mRNAs linked to cancer development, in melanoma. Understanding the spatiotemporal dynamics of these interactions *in vivo,* and a thorough knowledge of the interplay between RBPs and miRNAs is paramount to understanding the complexity of post-transcriptional gene control surrounding aberrant mRNA metabolism in skin cancers.

**Figure 6:**
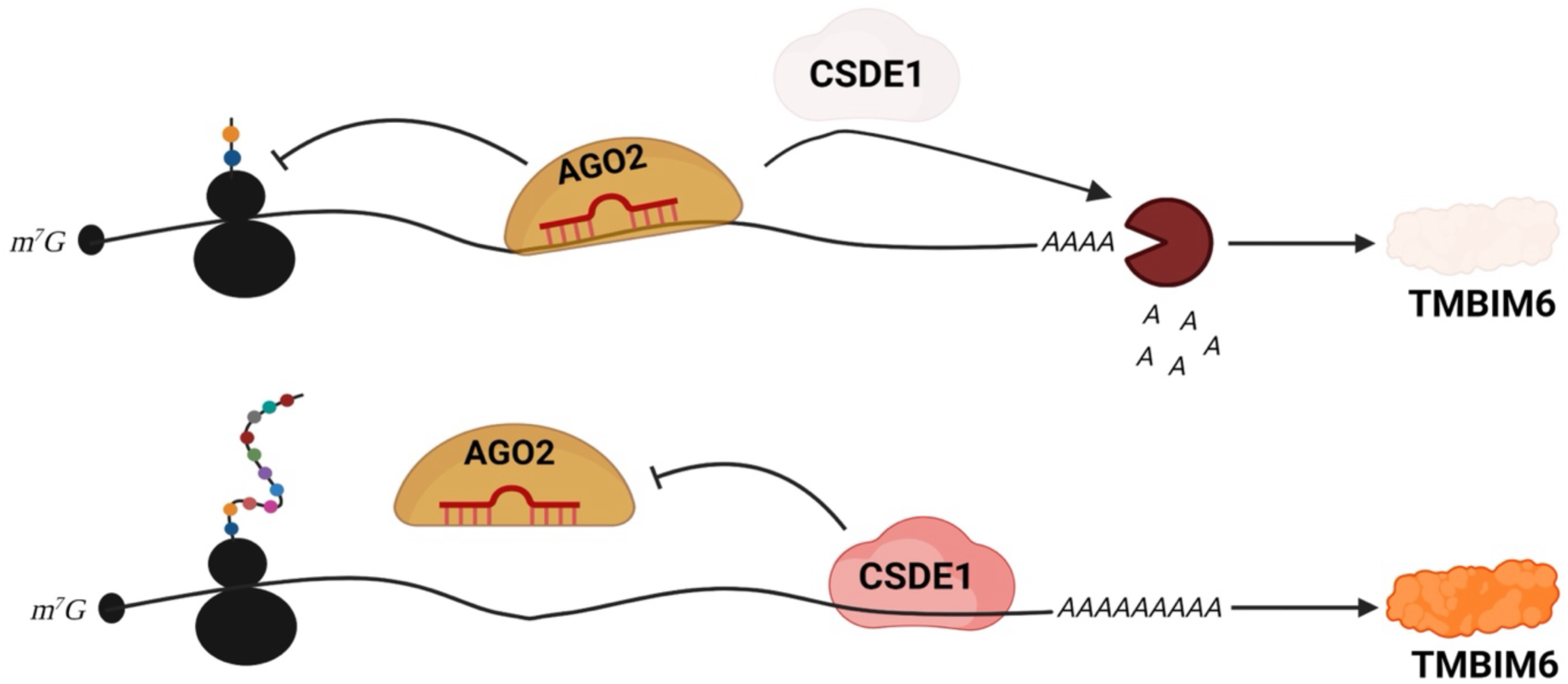
Illustration for the CSDE1-mediated attenuation of miR-20a-5p-guided silencing of TMBIM6. CSDE1 binds TMBIM6 mRNA and hinders miR-20a-5p/AGO2 interaction with the target mRNA, thus facilitating its de-repression.

## MATERIALS AND METHODS

### Cell Culture and transfections

The melanoma cell lines (WM-164, WM-793, 451-LU, 1205-LU, SK-Mel-147, SK-Mel-94, and UACC-62) and HEK293T cells were grown in Dulbecco’s modified Eagle’s medium (DMEM) (Sigma-Aldrich) supplemented with 10% fetal bovine serum (FBS) (Sigma-Aldrich), 50 U/ml penicillin and 50 μg/ml streptomycin (Sigma-Aldrich). The M-14 melanoma cell line was grown in RPMI with 10% fetal bovine serum, 50 U/ml penicillin, and 50 μg/ml streptomycin. Melanocytes were isolated from skin biopsy of normal human subjects and cultured as described previously (Goyer et al. 2019). Cells were grown in a humidified incubator at 37°C and 5% CO_2_. For transfections, the cell lines (UACC-62, SK-Mel-147) were seeded in 6-well plates or 100mm dishes at a confluency of 50%-80%, 24 hours before the transfection. miRNA mimic or antagomiR or siRNA transfections (50 nM) were performed using the jetPRIME transfection reagent (Polyplus). Plasmid DNAs were transfected using jetOPTIMUS transfection reagent (Polyplus) or X-tremeGENE™ HP DNA Transfection Reagent (Sigma-Aldrich), according to the manufacturer’s protocols. The cells were collected 48 hours post-transfection, for downstream experiments.

### Co-immunoprecipitation

Cells were lysed with 1X lysis buffer (25 mM Tris HCL pH 7.4, 150mM NaCl, 1% IGEPAL CA-630, 1mM EDTA and 5% glycerol) containing protease inhibitors (Sigma-Aldrich). The cell lysate was centrifuged at 15,000g for 15 minutes at 4°C, and the supernatant was collected and measured for protein concentration using Bradford reagent (Bio-Rad). Dynabeads Protein G (Thermo Fisher) (20ul for a total protein extract of 2mg and 10μg of antibody) were used to perform co-immunoprecipitation with specific antibodies, as previously described (Kakumani et al. 2020). Briefly, the cell lysate was precleared by incubating them with the Dynabeads at 4°C, following which, the lysate was transferred to an Eppendorf tube containing the antibody-bead complex, for incubation at room temperature (RT), for 90 minutes. Post-incubation, the beads were washed three times with 1X lysis buffer, and the samples were extracted either by adding SDS loading buffer for western blotting or TRIzol reagent (Sigma-Aldrich) for RNA extraction.

### 2′-O-methyl (2′-O-Me) pull-down

The cell lysate was prepared in 1X lysis buffer containing protease inhibitors and an RNase inhibitor (Thermos Fisher). The miRNA pulldown assay was performed as previously described (Kakumani et al. 2020). Briefly, the cell lysate was precleaned with M-280 Streptavidin Dynabeads (Thermo Fisher) cross-linked with the unrelated 2**′**-O-Me oligonucleotide probe (Luc), for 45 min at RT, on rotation. The supernatant was then incubated with the streptavidin beads conjugated with oligonucleotide probes complementary to specific miRNAs, for 45 min at RT, with gentle mixing on a rotator. The beads were washed three times with 1X lysis buffer, and the samples were extracted using the SDS loading buffer. In the case of RNase treatment, RNase solution was added to the beads and incubated at RT for 30 min. After the treatment, the beads were washed three times with 1X lysis buffer, and the samples were extracted by adding SDS loading buffer and heated to 98°C for 5 minutes.

Luc: 5’-Bio-CAUCACGUACGCGGAAUACUUCGAAAUGUC-3’

miR-20a-5p: 5’-Bio-UCUUCCUACCUGCACUAUAAGCACUUUAACCUU-3’

### Luciferase reporter assays

HEK293T cells were grown to 60–80% confluency in 24-well plates before the transfection. The reporter assays were conducted as previously described (Kakumani et al. 2021). Briefly, the control or the miRNA-specific mimics (50nM) were transfected with jetPRIME transfection reagent according to the manufacturer’s instructions. Twenty-four-hour post-transfection, the growth medium was replaced, and the cells were transfected with the empty psiCHECK2 vector or the recombinant plasmids containing either the wild-type TMBIM6 3’UTR or its mutant, using JetPRIME transfection reagent. The cells were lysed 48 hours post-transfection with 100 μl of 1X passive lysis buffer (Promega), and the luciferase activity was measured using the Dual-luciferase Reporter assay system (Promega). The luciferase reporter activity (renilla/firefly (RL/FL)) was measured using a multimode microplate reader (Bio Tek Synergy).

### Plasmids

The wild-type TMBIM6 3’UTR was cloned into XhoI and NotI sites in the psiCHECK2 vector. Its mutant, which disrupts the miR-20-5p binding site, was generated using a Q5 Site-Directed Mutagenesis kit (NEB), following the manufacturer’s instructions. The primers used are Forward: 5’-TTTACTGATGgcagaTAGTTTTTGGTCTGTTACCTG-3’, Reverse: 5’-ATGAACATTTATCTACTGCC-3’. The N-terminal FLAG-CSDE1 and its deletion mutants were cloned into BamHI and XhoI sites in the pCDNA vector as described previously (Kakumani et al. 2020). The point mutant of CSDE1, namely Y37A, was generated by site-directed mutagenesis using the primers Forward: 5’-CTTGACCTCTgccGGATTCATTC-3’ and Reverse: 5’-AGTTTTTCAATAACCCCAG-3’.

### Antibodies

Antibodies were acquired as follows: CSDE1 (Abcam, Cat# ab201688), Dicer (Santa Cruz Biotechnology, Cat# SC-136979; Novus Biologicals, Cat# NBP1-06520), AGO2 (Abcam, Cat# ab186733), β-actin (Abcam, Cat# ab49900), TMBIM6 (Proteintech, Cat# 26782-1-AP), CDK6 (Cell Signalling Technology, Cat# 3136T), PMEPA1 (Santa Cruz Biotechnology, Cat# sc-293372), FLAG (Sigma-Aldrich, Cat# F1804).

## ACKNOWLEDGEMENTS

We thank all the members of our laboratory for their helpful comments and suggestions.

## AUTHOR CONTRIBUTIONS

SR performed most of the experiments. TG analyzed protein expression across skin cancer cell lines. PKK conceived and supervised the study. SR wrote the manuscript with input from PKK and FG.

---

**Figure S1: CSDE1 does not affect the stability of TMBIM6 protein.**

**(A)** Western blotting of protein extracts from SK-Mel-147 cells transfected with CSDE1 siRNA and treated with MG132 at 10µM for 4 hours. The samples were run on SDS-PAGE, and TMBIM6, CSDE1, and CDK6 proteins were probed. β-actin served as a loading control. **(B)** Western blotting of lysates from different melanoma cells (Metastatic: SK-Mel-147, SK-Mel-94, UACC-62, M14, 451 LU, 1205-LU; Non-metastatic: WM-164, WM-793) and non-tumoral melanocytes. The proteins indicated were probed using respective antibodies. Vinculin was used as a loading control.

